# Tracking genetic invasions: genome-wide SNPs reveal the source of pyrethroid-resistant *Aedes aegypti* (yellow fever mosquito) incursions at international ports

**DOI:** 10.1101/490714

**Authors:** Thomas L. Schmidt, Anthony R. van Rooyen, Jessica Chung, Nancy M. Endersby-Harshman, Philippa C. Griffin, Angus Sly, Ary A. Hoffmann, Andrew R. Weeks

**Affiliations:** Bio21 Institute, School of BioSciences, The University of Melbourne, Victoria 3010, Australia; cesar Pty Ltd, 293 Royal Parade, Parkville Victoria 3052, Australia; Bio21 Institute, School of BioSciences, The University of Melbourne, Victoria 3010, Australia; Melbourne Bioinformatics, The University of Melbourne, Victoria 3010, Australia; Department of Agriculture and Water Resources, Qantas Drive, Brisbane Airport, Queensland 4008, Australia; Bio21 Institute, School of BioSciences, The University of Melbourne, Victoria 3010, Australia; cesar Pty Ltd, 293 Royal Parade, Parkville Victoria 3052, Australia

**Keywords:** genome-wide SNPs, biological invasions, *Aedes aegypti*, insecticide resistance, assignment tests, biosecurity, discriminant analysis of principal components (DAPC), invasion pathways

## Abstract

Biological invasions are increasing globally in number and extent despite efforts to restrict their spread. Knowledge of incursion pathways is necessary to prevent new invasions and to design effective biosecurity protocols at source and recipient locations. This study uses genome-wide single-nucleotide polymorphisms (SNPs) to determine the origin of 115 incursive *Aedes aegypti* (yellow fever mosquito) detected at international ports in Australia and New Zealand. We also genotyped mosquitoes at three point mutations in the voltage-sensitive sodium channel (*Vssc*) gene: V1016G, F1534C, and S989P. These mutations confer knockdown resistance to synthetic pyrethroid insecticides, widely used for controlling invertebrate pests. We first delineated reference populations using *Ae. aegypti* sampled from 15 locations in Asia, South America, Australia and the Pacific Islands. Incursives were assigned to these populations using discriminant analysis of principal components (DAPC) and an assignment test with a support vector machine predictive model. Bali, Indonesia, was the most common origin of *Ae. aegypti* detected in Australia, while *Ae. aegypti* detected in New Zealand originated from Pacific Islands such as Fiji. Most incursives had the same allelic genotype across the three *Vssc* gene point mutations, which confers strong resistance to synthetic pyrethroids, the only insecticide class used in current, widely-implemented aircraft disinsection protocols endorsed by the World Health Organisation (WHO). Additionally, all internationally-assigned *Ae. aegypti* had *Vssc* point mutations linked to pyrethroid resistance that are not found in Australian populations. These findings demonstrate that protocols for preventing introductions of invertebrates must consider insecticide resistance, and highlights the usefulness of genomic datasets for managing global biosecurity objectives.

## Introduction

Invasive alien species are a major threat to global biodiversity, agriculture, and human health, and confer a great economic burden upon developed and developing countries alike (IUCN, 2000). Invasive species have been shown to disrupt soil processes (Purnima, Raghubanshi, & Singh, 2008) and ecological community structure (Hejda, Pyšek, & Jarošík, 2009), and are linked to reductions in the abundance and richness of native species (Blackburn, Cassey, Duncan, Evans, & Gaston, 2004). They are responsible for the global spread of human diseases such as malaria, dengue, Zika, and chikungunya that together cause over 700,000 deaths annually and create heavy burdens on health systems worldwide (Lounibos, 2002). The spread of invasive species is closely tied to human economic activity, with many species transported as ‘stowaways’ on human trade and transport vessels (Levine & D’Antonio, 2003; Westphal, Browne, MacKinnon, & Noble, 2008). Following the expansion of trade and transport activity in recent decades, global distributions of invasive species have increased markedly (Hulme, 2009), with current predictions still arguably underestimating their true extent (McGeoch et al., 2010).

Preventing the establishment of new alien species requires knowledge of incursion pathways (IUCN, 2000). Although border interceptions can be effective at stopping incursions (Bacon, Bacher, & Aebi, 2012; Caley, Ingram, & De Barro, 2015), resource limitations restrict the number of inspections on arriving goods and conveyances or the number of border sections that can be monitored. These strategies must therefore seek to maximise detections by incorporating knowledge of incursion pathways, such as more frequent inspection of goods and conveyances from likely source locations (Robinson, Burgman, & Cannon, 2011), or regular inspection of known entry points of highly destructive species (Mehta, Haight, Homans, Polasky, & Venette, 2007; Myers, Simberloff, Kuris, & Carey, 2000). Incursion pathways can also be monitored to estimate parameters indicating the likelihood of future invasions, such as propagule pressure and opportunities for selection (Wilson, Dormontt, Prentis, Lowe, & Richardson, 2009).

Genomic information can be a useful means for identifying source populations of incursions. Invading individuals of unknown source intercepted at borders (hereafter defined as “incursives”) can be genotyped using genetic markers and compared with reference populations from the species’ known distribution. Analyses can then be performed to assign incursives to reference populations, in order to identify the likely source population or geographic region of origin. Genome-wide single nucleotide polymorphisms (SNPs) in particular can provide clear geographical delineation of populations (Rašić, Filipović, Weeks, & Hoffmann, 2014), which allows for confident assignment of intercepted individuals to their population or region of origin. This method can in theory enable the assignment of all incursive individuals, and thus improves upon previous methods that either rely on the interception of incursives at the border as they arrive from a known location (Mehta et al., 2007), or that use lower-resolution genetic markers that frequently cannot resolve individual origin to a fine geographic scale (Puckett & Eggert, 2016). Genome-wide SNPs could also be used to identify the source of incursions post-border and improve the chance of eradication by using pathway management to prevent future incursions.

*Aedes aegypti* (L.) (yellow fever mosquito) is a highly invasive pest and a primary global vector of arboviruses such as dengue (C. E. G. Smith, 1956), Zika (Bogoch et al., 2016) and chikungunya (Weaver & Lecuit, 2015). Its pantropical distribution has grown rapidly over the last century (Kraemer et al., 2015), placing a greater part of the human population at risk of disease. In Australia, *Ae. aegypti* is currently only endemic to Queensland, but once had a broader distribution that extended into New South Wales, South Australia and Western Australia (Russell et al., 2009). Under current climate change scenarios, there is speculation that the species will increase its distribution in Australia again (Kearney, Porter, Williams, Ritchie, & Hoffmann, 2009; Russell et al., 2009). Furthermore, detections of *Ae. aegypti* at Australia’s international airports increased markedly over the period from 2014 to 2016 (Fig 1), with 39 detections at Perth International Airport alone between October 2015 and April 2016. New Zealand does not currently support a local *Ae. aegypti* population, but historically has received incursives from places such as Fiji (Laird, 1951).

**Fig 1:**
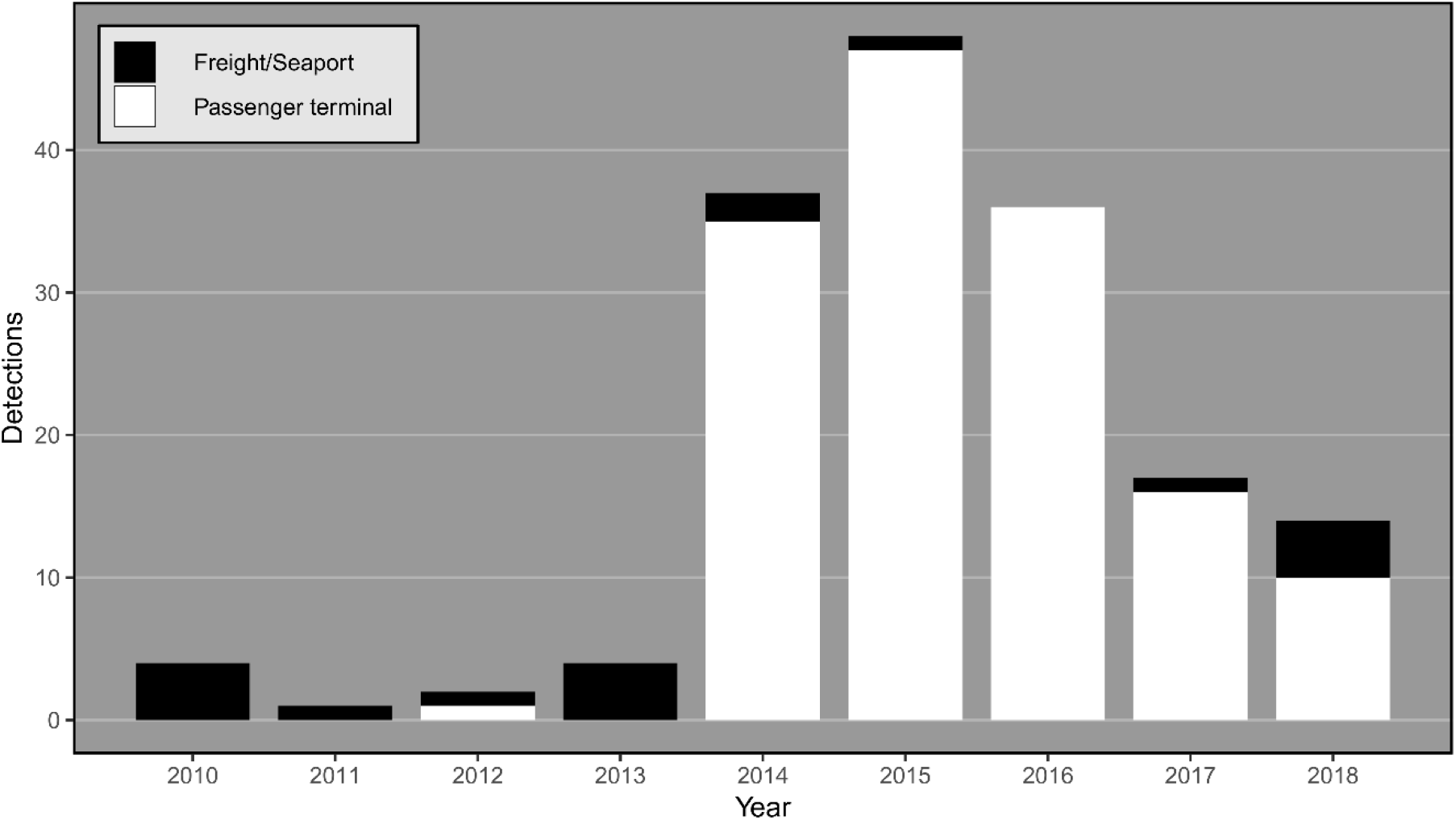
Number of *Ae. aegypti* detections per year at Australian international terminals. Detections at passenger airline terminals are shown in white, while those at seaports or freight terminals are in black. Each detection represents either a single adult retrieved from a trap or one or more larvae or pupae retrieved from an ovitrap.

When dispersing by flight alone, *Ae. aegypti* is a weak disperser that prefers to remain close to human dwellings and sites of activity (Harrington et al., 2005). Its current distribution has been facilitated by several centuries of human transport, including terrestrial (Guagliardo et al., 2014), marine (J. E. Brown et al., 2014) and aerial (Sukehiro et al., 2013) conveyance. Studies of *Ae. aegypti* using genome-wide SNPs have found genetic structure at fine scales of < 5 km (Schmidt, Filipović, Hoffmann, & Rašić, 2018) and strong population differentiation at broader scales (Rašić et al., 2015, 2014). Similar SNP datasets to that used in this study have proved useful for investigating the source of newly established *Ae. aegypti* populations (J. E. Brown et al., 2014; Gloria-Soria et al., 2018). The high level of genetic structuring at local and regional scales in this species suggests that genome-wide SNP analyses are likely to be informative for determining the origin of incursive mosquitoes, provided reference mosquitoes are available in sufficient numbers from a range of potential source populations.

The increasing rate of incursive *Ae. aegypti* detections at Australian airports has occurred despite protocols that ensure all international flights into Australia undergo disinsection procedures (DAWR/MPI, 2016), wherein insecticides are used to kill incursives through residual and/or knockdown treatments before or after take-off. However, dispersing pests from populations resistant to insecticides may be less affected by these measures. Although such resistance is not present in Australian *Ae. aegypti* (Endersby-Harshman et al., 2017), it is common in *Ae. aegypti* populations globally (Endersby-Harshman, Weeks, & Hoffmann, 2018; Ranson, Burhani, Lumjuan, & Black, 2010; L. B. Smith, Kasai, & Scott, 2016). Point mutations at the *Vssc* gene can confer target-site knockdown resistance to synthetic pyrethroids in a broad range of insects (Scott et al., 2013; Scott, Yoshimizu, & Kasai, 2015; Seong et al., 2010) including *Ae. aegypti* (Donnelly et al., 2009). The only two chemicals registered for aircraft disinsection by the World Health Organisation (WHO), permethrin and d-phenothrin (IPCS, 2013; WHO, 2012), are synthetic pyrethroids.

In this study, we use genome-wide SNPs to assign *Ae. aegypti* intercepted at international ports in Australia and New Zealand to their likely population, country, or region of origin, by comparing incursives with reference mosquitoes collected from 15 locations around the world. We also genotype incursive and reference mosquitoes for three point mutations at the *Vssc* gene known to confer resistance to synthetic pyrethroids in *Ae. aegypti* (Wuliandari et al., 2015). Our results are informative to the control and monitoring of a broad range of invasive invertebrates, and demonstrate how population genomic data can elucidate risks associated with species and genetic incursions.

## Materials and Methods

### Mosquito collection

A total of 115 incursive *Ae. aegypti* were collected from passenger and freight terminals at airports and at seaports and Approved Arrangement facilities in Australia and New Zealand between December 27^th^, 2012 and February 26^th^, 2018. Mosquitoes were considered to be incursive if they fit either of two conditions: i) they were collected at a terminal outside of Queensland, within which *Ae. aegypti* is endemic; or ii) they were collected within Queensland, but had alleles conferring resistance at the *Vssc* gene that are not normally found in Australian *Ae. aegypti* (Endersby-Harshman et al., 2017). Overall, only a single incursive mosquito from Queensland was included (collected from Townsville seaport); all other incursives were collected outside the native range. The majority of incursives were collected at international airport passenger terminals (73.0%), generally around baggage areas airside. Appendix A lists data for each of the 115 incursives.

For the reference populations, we used 18 samples of *Ae. aegypti* collected throughout its range, with emphasis placed on regions with high frequency of passenger air traffic to Australia and New Zealand (Fig 2). This included samples from 15 locations: Bali (Indonesia), Yogyakarta (Indonesia), Kuala Lumpur (Malaysia), Bangkok (Thailand), Rio de Janeiro (Brazil), Cairns (Australia), Townsville (Australia), Nha Trang (Vietnam), Ho Chi Minh City (Vietnam), Fiji, Vanuatu, Kiribati, New Caledonia, Taiwan and Singapore. Bali, Kuala Lumpur and Rio de Janeiro were each sampled twice at different time points to determine the stability of genetic patterns. Where possible, we sampled close to airports with the highest frequency of passenger traffic into Australia. In the case of Rio de Janeiro, from which there are no direct flights into either Australia or New Zealand, the samples were intended to represent a general South American or Pan-American reference population, with specific assignments within this continent prohibited without samples from other locations. Appendix B lists data for each of the 18 reference samples.

**Fig 2:**
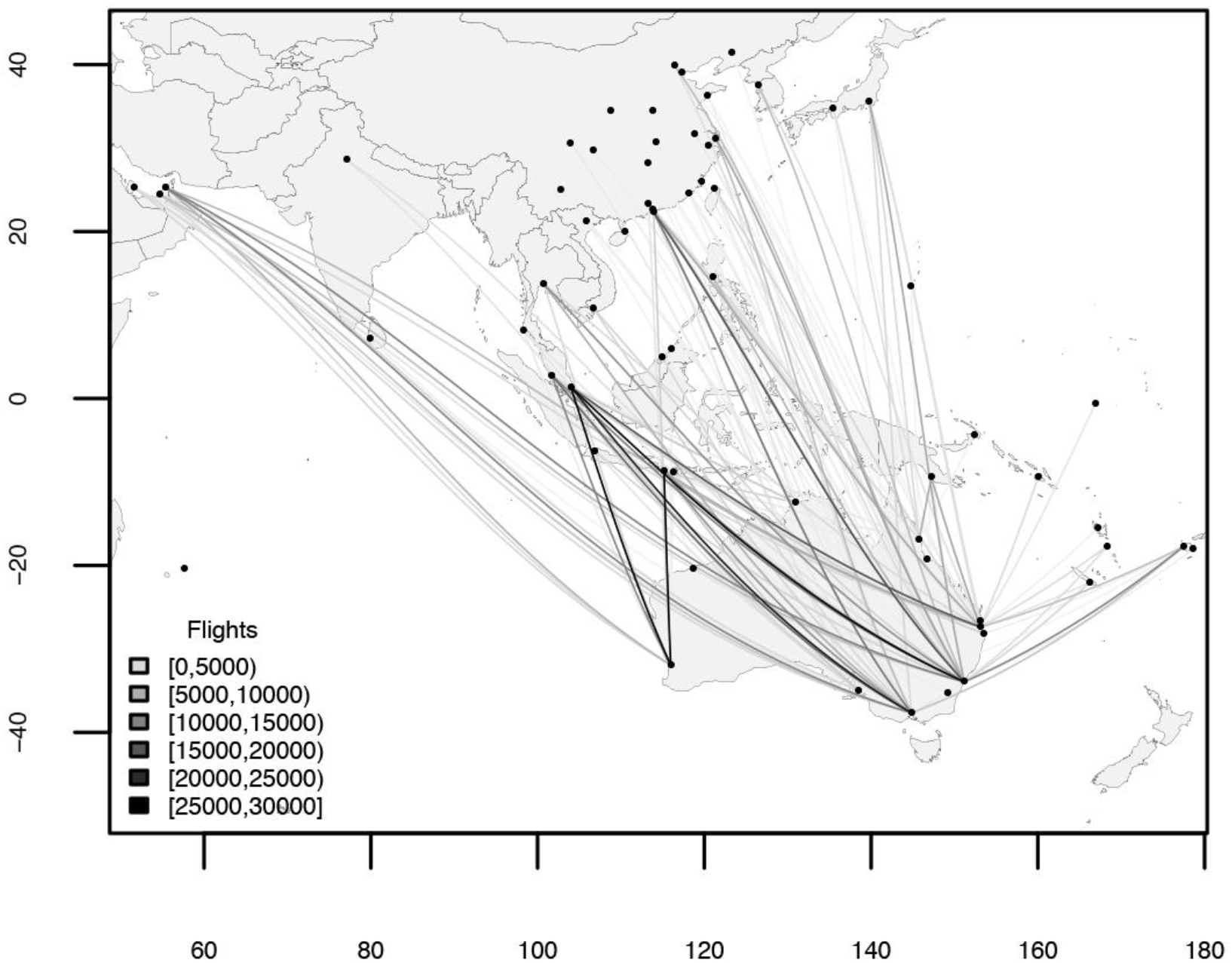
International passenger flights into Australia, 2014 – 2018. Total number of international passenger flights into Australian airports between January, 2014, and June, 2018, from regions in Asia, the South Pacific, and the Middle East.

DNA was extracted from mosquitoes using either Qiagen DNeasy Blood & Tissue Kits (Qiagen, Hilden, Germany) or Roche High Pure™ PCR Template Preparation Kits (Roche Molecular Systems, Inc., Pleasanton, CA, USA), each with an RNase A treatment step. Extracted DNA was used for SNP detection and genotyping at the *Vssc* gene.

### SNP detection and filtering

We applied the double digest restriction-site associated DNA sequencing (ddRADseq) protocol for *Ae. aegypti* developed by Rašić et al. (2014) to construct RAD libraries. The ddRADseq approach provides markers with excellent resolution at intraspecific scales (Peterson, Weber, Kay, Fisher, & Hoekstra, 2012), and is therefore suitable for both broad and fine scale population assignment in this species, as well as for deciphering patterns of pairwise relatedness to help optimise analyses of genetic structure. Libraries were sequenced at either the Australian Genome Research Facility (AGRF, Melbourne, Australia) or the University of Melbourne (Pathology) on a HiSeq 2500 (Illumina, California, USA) in either High Output or Rapid Run modes.

We used the process_radtags program in Stacks v2.0 (Catchen, Hohenlohe, Bassham, Amores, & Cresko, 2013) to demultiplex sequence reads and trim the reads to 80 bp in length. Using a 15 bp sliding window, low quality reads were discarded if the average phred score dropped below 20. Reads were aligned to the *Ae. aegypti* nuclear genome assembly AaegL4 (Dudchenko et al., 2017) with Bowtie v1.2.1.1 (Langmead, Trapnell, Pop, & Salzberg, 2009), allowing for a maximum of three mismatches (-v 3 --tryhard --best). The Stacks pipeline was used in three stages: initial filtering of reference mosquitoes; establishment of reference populations from the 18 reference samples; and building a final catalog of incursive and reference mosquitoes for population assignment.

Initial filtering of reference samples aimed to remove closely related mosquitoes and those with high levels of missing data, and to normalise the number of mosquitoes in each reference population. We used the Stacks *ref_map* pipeline to build individual Stacks catalogs for each of the 18 reference samples, from which we called genotypes at RAD stacks at a 0.05 significance level. We generated VCF files for each catalog with the Stacks program *populations*. SNPs were required to be present in ≥ 75% of the mosquitoes and have a minor allele frequency ≥ 0.05 (−r 0.75 --min_maf 0.05 --vcf). We used *VCFtools* (Danecek et al., 2011) to remove loci that deviated from Hardy-Weinberg equilibrium, then thinned the data so that no two SNPs were within 250 kbp of one another. As the *Ae. aegypti* genome contains approximately 2.1 Mbp sequence per centimorgan (S. E. Brown, Severson, Smith, & Knudson, 2001), thinning the data in this way retains up to 8 SNPs per centimorgan, a density shown to sufficiently filter out linked loci (Cho & Dupuis, 2009). Reference mosquitoes with ≥ 30% of missing genotypes were omitted from all further analyses. We used SPAGeDi (Hardy & Vekemans, 2002) to calculate Loiselle’s *k* (Loiselle, Sork, Nason, & Graham, 1995) among reference mosquitoes, identifying pairs with putative first-degree relatedness (*k* ≥ 0.1875; Iacchei et al., 2013) and omitting related mosquitoes in order of missing data so that all remaining pairs had *k* < 0.1875. Finally, for reference samples with more than 18 mosquitoes, we omitted mosquitoes in order of missing data to reduce the number to 18.

To establish reference populations from the reference samples, we generated a new catalog containing all of the filtered reference mosquitoes. We retained SNPs that were present in ≥ 75% of the mosquitoes in each reference sample and that had minor allele frequencies ≥ 0.05. We used the R package *assignPOP* v1.1.4 (Chen et al., 2018) to perform Monte-Carlo cross-validation on reference samples (functions “assign.MC” and “accuracy.MC”), treating all samples as putative populations. In this process, a proportion of mosquitoes was resampled from each putative population, and this subset of mosquitoes were assigned to populations using the remaining mosquitoes from each putative population. We ran this cross-validation multiple times, in turn resampling a proportion of 0.1, 0.2, and 0.3 of the mosquitoes from each reference sample, with 200 replicates run for each proportion.

Reference samples were treated as distinct reference populations if resampled mosquitoes were consistently assigned to their correct sample. In cases where resampled mosquitoes were assigned across multiple reference samples, we created composite reference populations from the multiple samples. If any composite reference populations contained more than 18 mosquitoes, we omitted mosquitoes in order of missing data proportion to reduce the number to 18.

A final catalog was built containing the 115 incursives and the 188 reference mosquitoes retained after filtering. This catalog was used to produce subsets of SNPs for the analyses delineated below. In each subset, we retained SNPs with minor allele frequency ≥ 0.05 that were present in ≥ 75% of the mosquitoes in each reference population and in ≥ 75% of incursives.

### Geographical assignment of incursive *Ae. aegypti*

Population assignment comprised two separate processes: cluster detection with discriminant analysis of principal components (DAPC; Jombart, Devillard, & Balloux, 2010), performed in the R package *adegenet* v2.1.1 (Jombart, 2008); and a Monte-Carlo assignment test using a support vector machine predictive model, performed in *assignPOP*. We used DAPC to partition reference mosquitoes and incursives into sets of clusters, and from these partitions infer population assignment. We used *assignPOP* to generate for each incursive a posterior probability of assignment to each of the reference populations, and from these probabilities infer confidence in assignment. Incursives showing inconsistent assignment for DAPC and *assignPOP* were not considered well-assigned.

For population assignment with DAPC, we used the *adegenet* function “find.clusters” to establish partitions within 115 subsets of mosquitoes. Each subset included all 188 reference mosquitoes and a single incursive. We identified an optimal range for the number of clusters (*K*) to use for partitioning, using 1,000 principal components and 10^9^ iterations for each *K*. The lower bound of this range was set as the value of *K* with the lowest Bayesian information criterion (BIC) and the upper bound was the value with the lowest Akaike information criterion (AIC), which provided a range of *K* from the conservative BIC to the less conservative AIC (Raftery, 1999). For each subset and for each value of *K* within this range, the single incursive formed a cluster containing the incursive and one or more of the reference populations. This clustering indicated the assignment of the incursive for that value of *K*. Following this, we performed cluster detection among closely related reference populations, which allowed the use of more SNPs and thus more confident assignment among populations within a subregion. For incursives initially assigned to Asian populations (Bali, Bangkok, Kuala Lumpur, Singapore, Taiwan, Vietnam, and Yogyakarta), we performed cluster detection using only the Asian reference populations, and did likewise for incursives initially assigned to the Pacific Islands (Fiji, Kiribati, New Caledonia, and Vanuatu).

For population assignment in *assignPOP*, we used the function “assign.X” to perform assignment tests on incursives. This analysis used the 188 reference mosquitoes to build a predictive model with which to assign the 115 incursives, using all principal components with eigenvalues > 1, and a linear support vector machine classification function. This generated posterior probabilities of membership for each individual to each reference population. From each of these scores we calculated ‘relative probability’ of membership, which was defined as the probability of membership to the most likely population divided by the probability of membership to the second most likely population. We used these relative probabilities as an additional measure of confidence in assignment. For an incursive to be considered well-assigned it would need to have a relative probability > 2; that is, the posterior probability of its assigned reference population would have to be at least twice that of the next most likely population.

### *Vssc* Genotyping

All incursives were genotyped for the presence of three target site mutations (V1016G, F1534C, S989P) in the *Vssc* gene known to provide resistance to synthetic pyrethroids (Du et al., 2013; Wuliandari et al., 2015). We also genotyped mosquitoes from the Bali and Fiji reference populations. The mutation site amino acid positions in this study are labelled according to the sequence of the most abundant splice variant of the house fly, *Musca domestica, Vssc* (GenBank accession nos. AAB47604 and AAB47605) (Kasai et al., 2014). The V1016G mutation is widespread in Southeast Asia and is known to provide resistance to Type I and Type II synthetic pyrethroids when in the homozygous (GG) state (Wuliandari et al., 2015). The F1534C mutation is thought to confer resistance to Type I synthetic pyrethroids (Du et al., 2013) when homozygous. The S989P mutation alone has no effect, but in the presence of the V1016G mutation it is thought to act synergistically, increasing resistance to Type II synthetic pyrethroids (but not Type I synthetic pyrethroids) (Hirata et al., 2014). We developed TaqMan^®^ assays for all three target site mutations based on the high resolution melt genotyping assays in Wuliandari et al. (2015). Genotyping of each SNP was then undertaken on a LightCycler^®^ 480 (Roche, Basel, Switzerland) 480 real time PCR machine with three replicates in a 384-well plate format.

## Results

### Delineation of reference groupings

Monte-Carlo cross validation using *assignPOP* showed perfect assignment of resampled mosquitoes among 8 of the 18 reference samples, indicating that these samples were genetically distinct and could be treated as separate reference populations. However, for the time-separated pairs of reference samples from Bali, Kuala Lumpur, and Rio de Janeiro, resampled mosquitoes were frequently assigned to the incorrect sample of the pair. This indicates that geographical genetic structure observed at these locations was stable over time. Additionally, resampled mosquitoes from the Cairns and Townsville reference samples showed 10% incorrect assignment with each other, which is not surprising given weak genetic structure across this area (Endersby et al., 2011), while 7% of resampled mosquitoes from the Ho Chi Minh City and Nha Trang reference samples were incorrectly assigned. Accordingly, each of these five pairs were combined into composite reference populations, leaving 13 reference populations for the assignment of incursives.

### Geographical assignment of incursive *Ae. aegypti*

DAPC cluster detection using 41,834 SNPs gave a range for the number of clusters (*K*) of 3 ≤ *K* ≤ 13. At *K* = 3, populations were partitioned into a South American cluster, an Asian cluster, and a cluster containing Australia and the Pacific Islands. The Australian population split from the Pacific Islands at *K* = 4, following which new geographical partitions formed at each increase until *K* = 13 placed all reference populations into their own cluster. These same geographical partitions were established at *K* = 13 in each run, indicating that the inclusion of the single incursive was not biasing cluster detection.

Of the 115 incursives, 111 demonstrated clustering for 3 ≤ *K* ≤ 12 that was consistent with their final cluster assignment at *K* = 13 (Appendix A). The 8 incursives collected at New Zealand terminals all showed consistent clustering with the Pacific Islands, while the incursives collected in Australia clustered with Southeast Asia, Australia, and South America. Supplementary Fig A shows a typical stepwise delineation of clusters for 3 ≤ *K* ≤ 13 for incursive IPT33.

Posterior probabilities of membership to the 13 reference populations calculated for incursives using *assignPOP* were of range 0.16 – 0.82. Relative probabilities (see Materials and Methods) ranged from 1.01 to 29.00. For incursives consistently assigned to either of the Asian or Pacific Islands clusters, we repeated assignment tests using only those reference populations, which allowed us to increase confidence in assignment by incorporating more SNPs: 55,443 and 146,142 respectively. Overall, 92 incursives (80%) had relative probabilities > 2, while 23 (20%) had relative probabilities < 2. Comparing assignment between DAPC and *assignPOP*, 112 of the 115 incursives (97.4%) had highest posterior probabilities for the same reference population they were assigned to at *K* = 13 (Appendix A), indicating strong agreement between the two assignment methods.

Fig 3 shows the relationship between posterior probability, relative probability, and proportion of missing data. The incursives with relative probabilities > 2 (white and black circles) formed a group in which posterior probability of membership decreased as missing data increased. Five incursives with relative probabilities > 2 were exceptions to this (black circles), grouping with the incursives with relative probabilities < 2 (black squares). This latter group (black circles and squares) had low posterior probabilities and mostly low missing data, and these variables were not well correlated (linear regression: R^2^ = 0.023, F_27_ = 0.609, P = 0.442). Among the incursives of the first group (white circles), there was a strong negative correlation between posterior probability of membership and proportion of missing data (linear regression: R^2^ = 0.795, F_86_ = 329.6, P < 0.001), indicating that lower posterior probabilities in this group were likely due to missing data (e.g. from imperfect preservation of specimens). Conversely, 25 of the 28 incursives in the second group (black circles and squares) had posterior probabilities < 0.5 despite having < 0.13 missing data (Fig 3). As the low posterior and relative probabilities of these incursives could not be explained by high missing data, it seems likely that these incursives were from source populations not sampled in this study.

**Fig 3:**
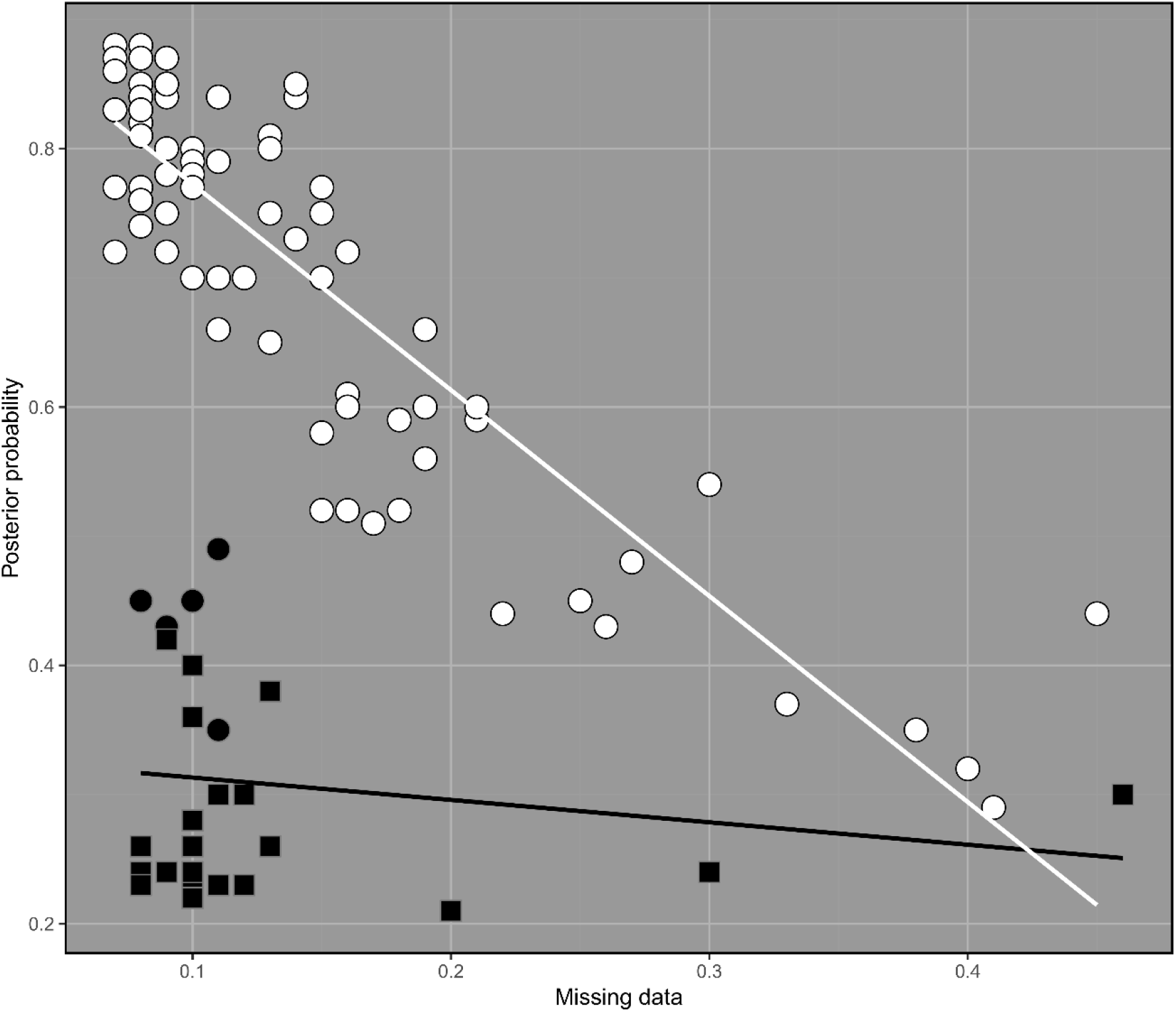
Posterior probabilities and proportion of missing data for the 115 incursive *Ae. aegypti*. White and black circles indicate incursives with relative probabilities > 2, while black squares indicate incursives with relative probabilities < 2. Incursives marked in white were considered well-assigned, while those marked in black were considered poorly-assigned. Among well-assigned incursives, lower posterior probabilities correlated strongly with higher missing data (linear regression: R^2^ = 0.795, F_86_ = 329.6, P < 0.001). Among poorly-assigned incursives there was no similar relationship (linear regression: R^2^ = 0.023, F_27_ = 0.609, P = 0.442).

We considered the 87 incursives (75.7%) represented as white circles in Fig 3 as well-assigned. These all showed consistent assignment for DAPC and *assignPOP* and consistent cluster assignment for 3 ≤ *K* ≤ 13, and had relative probabilities > 2. Fig 4 shows a DAPC of the well-assigned incursives plotted with the 13 reference populations. The well-assigned incursives were assigned to the following source populations: 62 to Bali, 13 to Kuala Lumpur, 7 to Rio de Janeiro, 3 to Australia, 1 to Fiji and 1 to Bangkok (Appendix A). The incursive assigned to Fiji was detected in New Zealand and was the only well-assigned New Zealand incursive; the incursives assigned elsewhere were all detected in Australia. Supplementary Fig B shows a DAPC of the seven Asian reference populations (Bali, Bangkok, Kuala Lumpur, Singapore, Taiwan, Vietnam, Yogyakarta) plotted with the 76 well-assigned incursives putatively from Asia.

**Fig 4:**
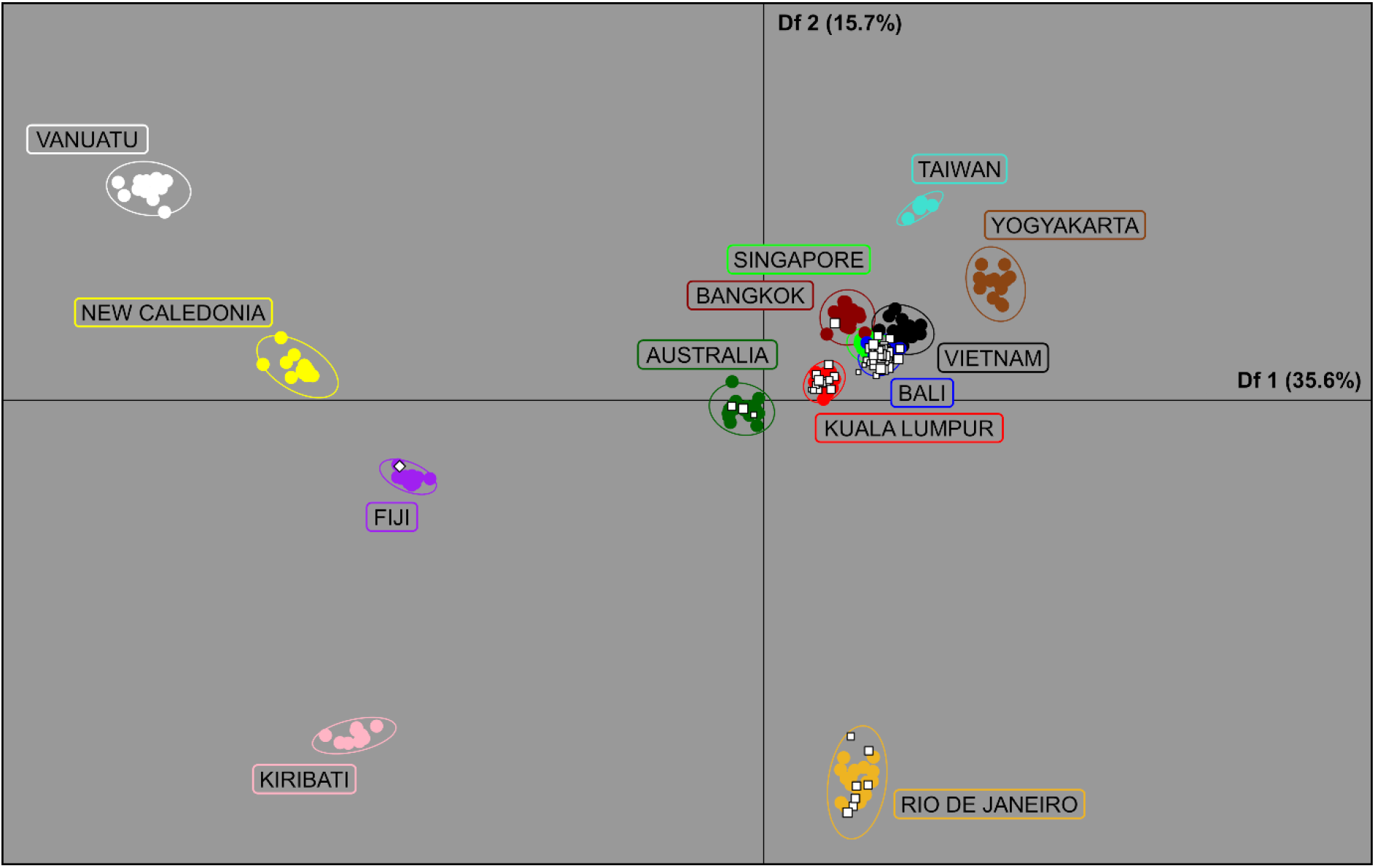
Discriminant analysis of principal components (DAPC). The plot shows the first and second discriminant functions of a DAPC of all 188 reference *Ae. aegypti* and all 87 well-assigned incursive *Ae. aegypti*, using 41,834 SNPs and 100 principal components. White squares in normal orientation show incursives intercepted at Australian terminals, while white squares rotated 45° show incursives intercepted at New Zealand terminals. Squares are sized by a logarithmic function indicating relative probability of membership (see Materials and Methods).

### *Vssc* Genotyping

TaqMan^®^ (Life Technologies, California, USA) assays for the three target site mutations showed that the most common genotype found in incursive mosquitoes across the three loci was homozygous resistant at V1016G, homozygous susceptible at F1534C, homozygous resistant at S989P (GG/TT/CC – the letters refer to the mutant or wildtype bases at each of the loci, rather than the amino acid notation). This genotype putatively confers strong resistance to Type I and Type II synthetic pyrethroids (Wuliandari et al., 2015). Of the 72 incursives with the GG/TT/CC genotype, 62 were assigned to Bali, 5 were assigned to Kuala Lumpur, 1 was assigned to Bangkok, and 4 were uncertain but had a likely Asian origin. As well as occurring in 100% of putatively Balinese incursives, this genotype was also found in all reference mosquitoes from Bali.

Another common genotype among incursives was homozygous susceptible at V1016G, homozygous resistant at F1534C, homozygous susceptible at S989P (TT/GG/TT), which putatively confers strong resistance to Type I synthetic pyrethroids (Du et al., 2013). This genotype was found in 18 incursive *Ae. aegypti* and 9 of the 11 Fijian reference mosquitoes. Of these incursives, seven were assigned to Rio de Janeiro, and one each to Bangkok and Fiji; the remainder were uncertain. Overall, 95 incursives (82.6%) had genotypes putatively conferring strong resistance to Type I synthetic pyrethroids.

Among all incursives, only three had the genotype that lacks all three resistance-associated point mutations (TT/TT/TT). These incursives were all well-assigned to the Australian reference population, indicating that they were likely transported via domestic means rather than an international pathway. This genotype was also found in one of the Fijian reference mosquitoes.

## Discussion

Assignment tests using genome-wide SNPs were able to identify the global source populations from which pyrethroid-resistant *Ae. aegypti* disperse into Australia and New Zealand. We observed evidence for different dispersal pathways into Australia and into New Zealand, as all the incursives intercepted in Australia were assigned to Southeast Asia, South America, and Australia, while all the incursives intercepted in New Zealand were assigned to the Pacific Islands. Among well-assigned incursives, most (71.2%) were assigned to Bali. All incursives assigned to Bali were collected airside at international passenger terminals in Australia, with the majority detected in Perth (56.3%), one of the most active routes between Australia and Denpasar International Airport in Bali (Fig 2). All *Ae. aegypti* suspected (incursives) and known (reference mosquitoes) to be from Bali shared a *Vssc* genotype (1016/1534/989; GG/TT/CC) linked to strong resistance to Type I and Type II synthetic pyrethroids (Wuliandari et al., 2015). Most of the suspected and known Fijian *Ae. aegypti* shared a genotype (TT/GG/TT) linked to strong resistance to Type I synthetic pyrethroids (Du et al., 2013). If the resistance observed in laboratory studies is indicative of field resistance, then the WHO aircraft disinsection protocols that rely on the use of Type I synthetic pyrethroids (IPCS, 2013; WHO, 2012) are unlikely to stop the dispersal of *Ae. aegypti* from regions where resistance to these insecticides is common. The protocols used in this study may be implementable for *Ae. aegypti* and other invasive species internationally, which may help indicate the extent to which pyrethroid-based protocols alone are insufficient for achieving global biosecurity objectives.

Our results strongly suggest that resistance to synthetic pyrethroids used during aircraft disinsection is likely to be contributing to the detections of *Ae. aegypti* at the Australian and New Zealand borders. There are currently only two chemicals registered for aircraft disinsection by the WHO, permethrin and d-phenothrin (IPCS, 2013; WHO, 2012); both are Type I synthetic pyrethroids (permethrin is first generation, d-phenothrin is second generation). Resistance to these pyrethroids varies substantially throughout populations of *Ae. aegpyti* worldwide (L. B. Smith et al., 2016), though Australian populations remain wholly susceptible (Endersby-Harshman et al., 2017). Of the 115 incursive mosquitoes analysed in this study, only three incursives had the wholly-susceptible genotype (TT/TT/TT). These were collected in Brisbane, Australia, and were assigned to the Australian population, and thus are likely to have been transported on a domestic transport route where disinsection was not conducted as required on all international flights to Australia (DAWR/MPI, 2016). Within most of the non-Australian *Ae. aegypti* reference populations in this study there is considerable diversity in resistance genotypes (Endersby-Harshman et al., 2018; unpublished data), with populations made up of homozygotes with theoretically strong resistance, susceptible wild-type homozygotes, and heterozygotes of varying resistance status. Our finding that all incursive mosquitoes from international sources had alleles associated with resistance to Type I pyrethroids, and that 82.6% were likely to have displayed strong levels of resistance, suggests that aircraft disinsection may be successfully eliminating pyrethroid-susceptible mosquitoes, while pyrethroid resistance may be permitting others to survive and reach the Australian and New Zealand borders.

International incursions of pyrethroid-resistant *Ae. aegypti* into Australia present three main management challenges. They could introduce alleles conferring pyrethroid resistance into the current, pyrethroid-susceptible Australian population and thereby reduce the effectiveness of current control programs; they could bring disease into non-endemic regions (Siala et al., 2015); and they could lead to the establishment of new Australian *Ae. aegypti* populations outside the current native range in Queensland. Considering the current high frequency at which resistant *Ae. aegypti* are being detected at the Australian border, it is critical that future research focuses on determining whether the target site mutations provide ‘operational resistance’ to disinsection and whether this differs for the various methods used for disinsection (e.g. pre-flight and top-of-descent versus pre-embarkation/residual treatment; DAWR/MPI, 2016). If operational resistance is shown to disinsection, then management of *Ae. aegypti* may need to focus more on suppressing populations at source locations. Interestingly, there has been a large reduction in the number of *Ae. aegypti* incursives detected at Australian airports over the last two years (Fig 1), which coincides with discussions between the Australian and Indonesian Governments as the results of this study were becoming available and may indicate a change in density of local mosquito populations in Bali either through local control efforts or other local dynamics. Nevertheless, due to the likely high incidence of pyrethroid resistance in these populations, it may be more effective to use non-insecticidal methods to suppress them, such as by releasing *Ae. aegypti* males that are sterile or that carry an introduced infection of endosymbiotic bacteria (McGraw & O’Neill, 2013). Local suppression around Denpasar International Airport may also help reduce the dengue burden in this area, which has one of the highest incidences of dengue in Bali (Purnama & Baskoro, 2013). Non-insecticidal suppression may also be useful for eliminating small populations of resistant mosquitoes in newly-invaded regions.

This study has applied protocols for the assignment of incursive individuals to their likely population of origin. Following their application to *Ae. aegypti*, our protocols could be applied to other invasive species that are also frequently detected at borders. One candidate would be the related dengue vector *Ae. albopictus*, which has invaded and continues to invade tropical and temperate regions globally (Benedict, Levine, Hawley, & Lounibos, 2007; Kraemer et al., 2015), including the Torres Strait Islands in northern Australia (Ritchie et al., 2006), from which it threatens invasion of the mainland. This approach may be limited by the weak genetic structure between *Ae. albopictus* populations compared with *Ae. aegypti* (Goubert, Minard, Vieira, & Boulesteix, 2016), even when investigated with high-density markers (Schmidt et al., 2017). However, genome-level population genetic studies on *Ae. albopictus* are currently scarce, and a large database of reference samples may be necessary to identify distinct populations and produce confident assignments as in this study.

For invasive species with population differentiation equivalent to or greater than that of *Ae. aegypti*, the methods used in this study are likely to provide similar sensitivity for determining the origin of incursions. Our methodology could also be applied for post-border detections, where newly invasive species are detected and there is a need to determine their likely source. For instance, *Diuraphis noxia* (Russian wheat aphid) was first detected in Australia in 2016, and successful control of this species now depends, in part, on identifying past and present sources of incursions (Yazdani et al., 2018). Although a recent genome assembly of *D. noxia* suggests that this species has low genetic variability (Burger & Botha, 2017), genome-wide SNPs may yet provide sufficient power to determine broad patterns of gene flow. Likewise, similar methodologies may help untangle the complexity of marine invasion narratives, in which many processes remain cryptic when analysed without the appropriate molecular tools (Ojaveer et al., 2014).

## Conclusions

Current WHO protocols for preventing the unintentional dispersal of invasive invertebrates rely upon Type I synthetic pyrethroids for disinsection of aircraft. This study demonstrates that these measures are likely to have been insufficient for preventing the dispersal of resistant strains of the major disease vector, *Ae. aegypti*, to the Australian and New Zealand borders through international air traffic. Most *Ae. aegypti* collected in Australia came from Bali, while all those collected in New Zealand came from Pacific Islands like Fiji. A large majority of incursives (82.6%) had point mutations at the *Vssc* gene that putatively confer strong resistance to Type I or Type I and Type II synthetic pyrethroids. These mutations were found at high frequency in the greater Balinese and Fijian *Ae. aegypti* populations. Our findings indicate that insecticide-based strategies are likely to be limited in their capacity to stop biological invasions when insecticide resistance is present, and alternative control strategies should be considered, particularly for important disease vectors such as *Ae. aegypti*.

## Data Archiving Statement

Aligned .bam files for 188 reference and 115 incursive *Ae. aegypti* have been archived at NCBI Genbank, accessible with accession number PRJNA522930.

## Appendix A

**Relevant data for each of the 115 incursives.** (uploaded separately)

## Appendix B

**Relevant data for each of the 18 reference samples.** (uploaded separately)

---

**Supplementary Fig A:**
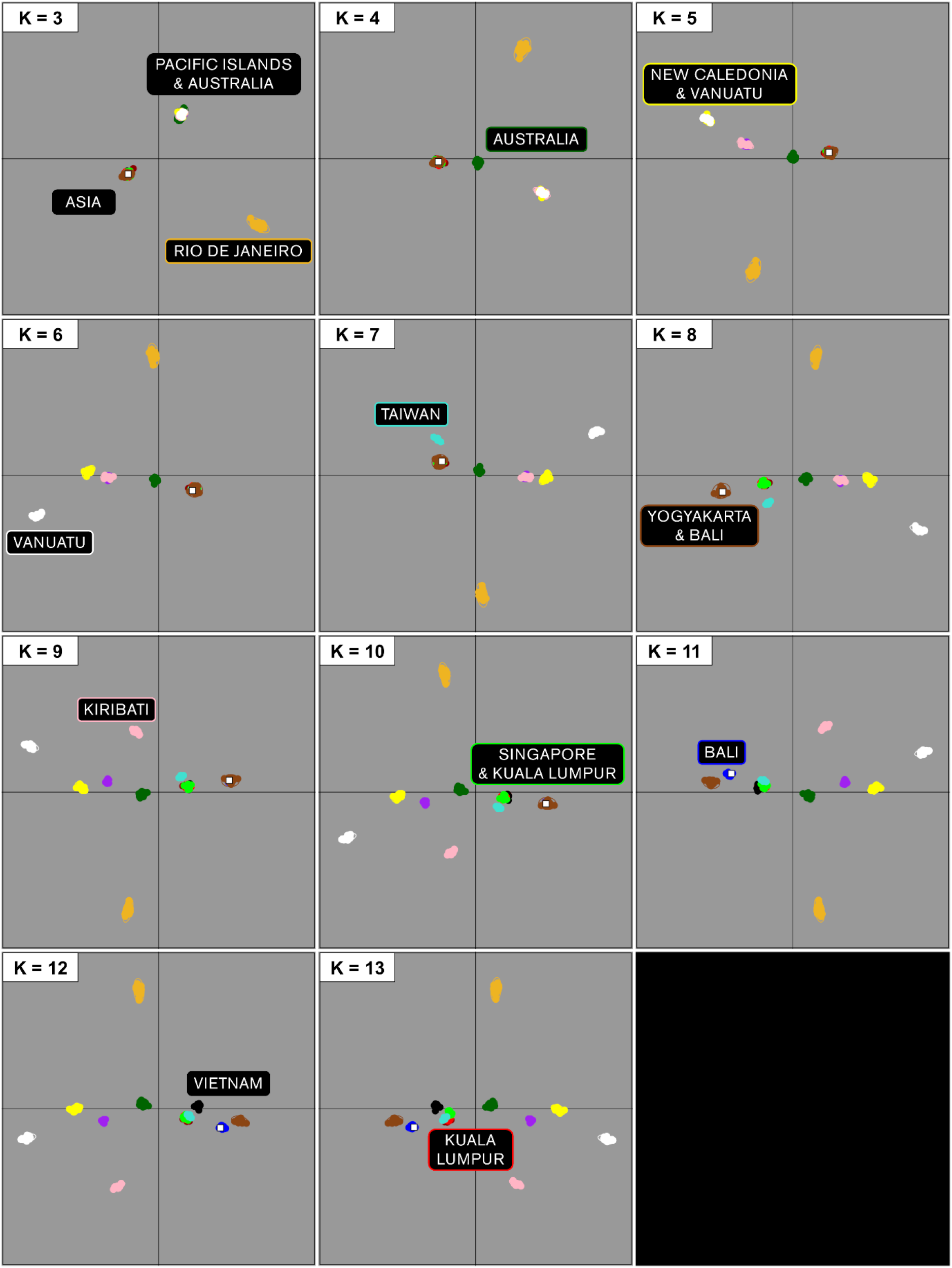
A typical stepwise delineation of clusters for 3 ≤ *K* ≤ 13 for incursive IPT33.

**Supplementary Fig B:**
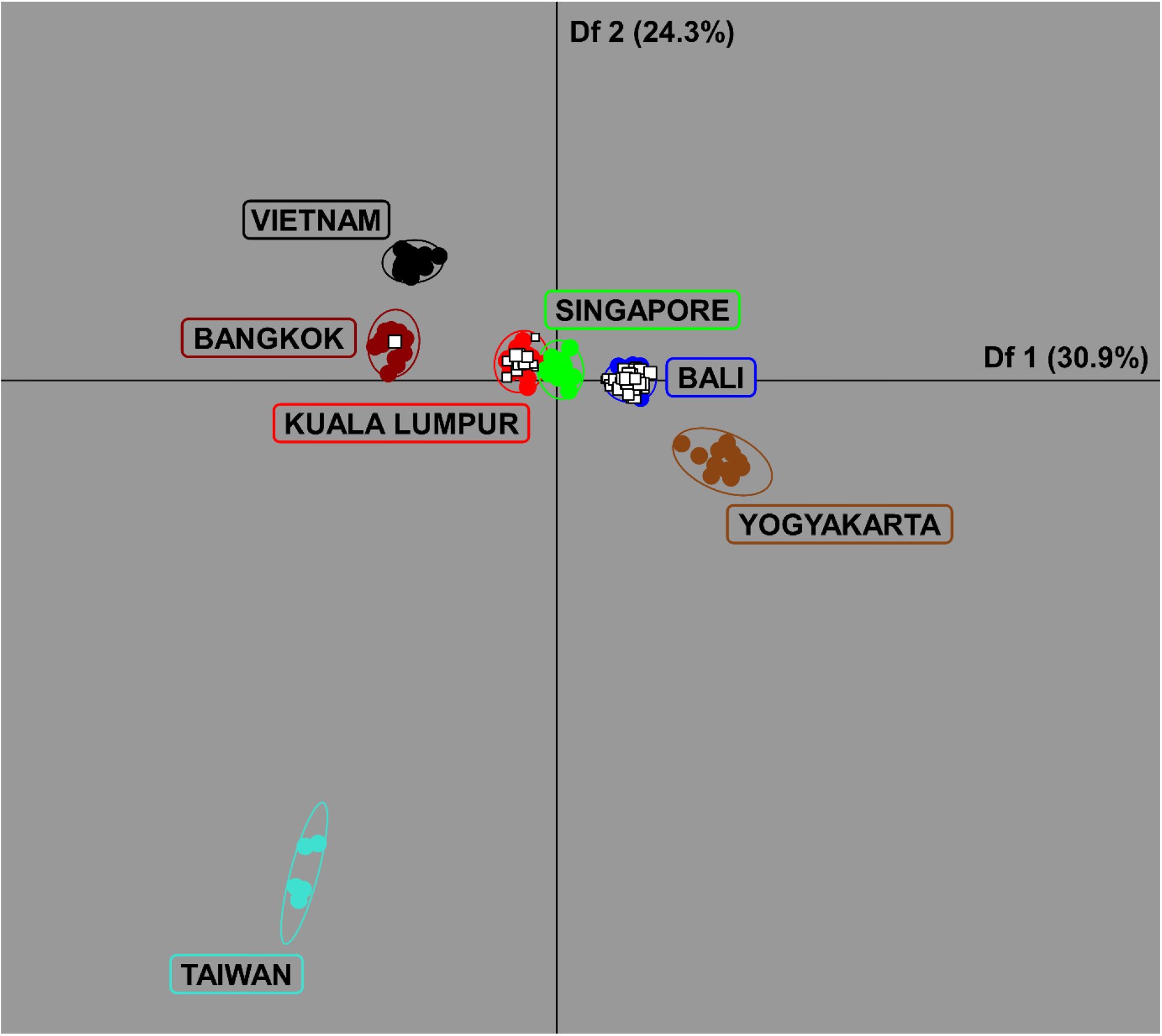
DAPC of the Asian reference populations and the 76 well-assigned incursives putatively from Asia.

## Acknowledgements

We thank Ashley Callahan, Jason Axford, Tim Hurst, Stephen Doggett, Elizabeth Valerie, Craig Williams and Joe Davis for the collection of reference samples used in this study. We also thank Ashley Callahan for conducting some of the DNA extraction and library preparation. We thank Gordana Rašić for input in conceptualisation and study design. We thank staff from the Department of Agriculture and Water Resources (DAWR), Australian Government, for the collection of all incursive mosquitoes and DAWR for providing funding for this work. AAH was supported by Program and Fellowship grants from the National Health and Medical Research Council (NHMRC), no. 1037003. TLS and AAH were also supported by the Wellcome Trust UK, no. 108508. These grants provided the necessary support to establish the reference database used in this study.

